# Correlative light and electron microscopy reveals fork-shaped structures at actin entry sites of focal adhesions

**DOI:** 10.1101/2022.04.27.489664

**Authors:** Karin Legerstee, Jason Sueters, Gert-Jan Kremers, Jacob P. Hoogenboom, Adriaan B Houtsmuller

## Abstract

Focal adhesions (FAs) are the main cellular structures to link the intracellular cytoskeleton to the extracellular matrix. FAs mediate cell adhesion, are important for cell migration and are involved in many (patho)-physiological processes. Here we examined FAs and their associated actin fibres using correlative fluorescence and scanning electron microscopy (SEM). We used fluorescence images of cells expressing paxillin-GFP to define the boundaries of FA complexes in SEM images, without using SEM contrast enhancing stains. We observed that SEM contrast was increased around the actin fibre entry site in 98% of FAs, indicating increases in protein density and possibly also phosphorylation levels in this area. In nearly three quarters of the FAs, these nanostructures had a fork shape, with the actin forming the stem and the high contrast FA areas the fork. In conclusion, the combination of fluorescent and electron microscopy allowed accurate localisation of a highly abundant, novel fork structure at the FA-actin interface.

## Introduction

Focal adhesions (FAs) are the main cellular structures to link the intracellular cytoskeleton to the extracellular matrix (ECM). They are typically several square micrometres in size^1,2^. On the membrane-facing side the main FA components are integrins, transmembrane receptors which directly bind to the extracellular matrix (ECM). F-actin, or filamentous actin, forms the edge of the FA on the cytoplasm-facing side. In between integrins and actin a large and diverse intracellular macromolecular protein assembly is present, for which over 200 different proteins have been reported^3,4^. These include (trans)membrane receptors, other than integrins, adaptor proteins and many different signalling proteins such as kinases, phosphatases and G-protein regulators, which through post-translational modifications add significantly to FA complexity. FAs experience force, the strength of which depends on the combination of myosin-II contractility and the stiffness of the ECM. Because of their importance in cell adhesion and to the transmission of force from the cell to the extracellular matrix, FAs are crucial to most types of cell migration, including in vitro over a 2D-surface. Migration and adhesion are key cellular functions required for many physiological and pathophysiological processes, like embryological development, the functioning of the immune system and cancer, in particular metastasis^4–6^.

The F-actin associated with FAs takes the shape of stress-fibres, a specialised form of actin associated with contractile myosin II and cytoskeletal proteins such as α-actinin^7^. There are two types of stress fibres associated with FAs: ventral stress fibres are associated with FAs at either end and typically transverse the whole cell, while dorsal stress fibres are linked to FAs on one end, typically near the cell front, then stretch upwards to the nucleus and the dorsal cell surface^8^.

Here we examine FAs and their associated F-actin fibres using a correlative fluorescence microscopy and Scanning Electron Microscopy (SEM) approach. Because FAs are very dense protein complexes found at the very edge of the cell and directly attached to the ECM, they are well suited to studying with SEM. We used cells stably expressing a fluorescently tagged form of the major FA protein paxillin to mark the FAs and fluorescently tagged phalloidin to stain the F-actin network. Although FAs have frequently been visualised using SEM^9–15^, overlaying the SEM images with the fluorescence images adds the possibility to clearly mark the FA boundaries in the SEM images. This revealed that FAs have a higher contrast in these images at the tip where actin fibres enter. Further examination revealed that these high contrasting FA areas and the associated F-actin fibre together have a forked shape, with the actin forming the stem and the high contrast areas within the FA forming the fork. Since no contrast-enhancing staining agent was applied this shows that protein density and possibly also phosphorylation levels are increased at the fork.

## Results and discussion

Here we examine FAs and their associated F-actin fibres using correlative fluorescence and scanning electron microscopy. Indium-Tin-Oxide (ITO) coated glass coverslips were additionally coated with a thin layer of type-I collagen to mimic the ECM, providing a surface for the integrin receptors to bind to. The ITO-coated glass provides a conductive substrate, which allows SEM inspection of thin samples such as cultured cells without metal shadowing or other conductive coating and/or without additional staining^16–21^. Onto these coverslips human bone cancer (U2OS) cells were seeded, stably expressing GFP-tagged paxillin to visualise the FAs. The cells were given 36 hours to adhere and form clear FAs, followed by chemical fixation (4% paraformaldehyde), permeabilization and staining with phalloidin to fluorescently stain the F-actin fibres. Immediately before imaging, the samples were dehydrated (ethanol). During imaging we were able to switch between fluorescent and SEM imaging without moving the stage^17^. However, in vacuum, under these dehydrated, permeabilised conditions, the GFP signal was relatively weak^18,22^. Therefore we first made the fluorescent images then followed up with the SEM images. To create overlay images the two images were scaled to the same size followed by a manual overlay procedure, mainly on the basis of the F-actin network as due to the phalloidin staining this is clearly visible in both imaging modalities. Of the 122 FAs with associated actin fibres, 97 were clearly visible in both modalities and the remaining 25 were unclear in the SEM images. These latter FAs were located further from the edge of the adherent part of the plasma membrane, where the cell is thicker. In these areas the FAs were often lost in the signal from the rest of the cell including other fibre networks, as no specific markers or contrast agents were used and the SEM at the low energies used only penetrates the upper tens of nm of the sample.

FAs are visible in the SEM images but the overlay images are needed to clearly define the boundaries of the FAs (Fig. 1). Especially where the FA connects to its F-actin fibre it is almost impossible to determine from the SEM-images alone where the F-actin fibre ends and the FA begins. However, using the GFP-paxillin signal to mark the FA boundaries, it becomes clear that in the SEM images the contrast of the FA complex is increased around the entrance point of the F-actin fibre. Note the SEM images show an apparent reverse contrast due to the comparatively high backscatter electron yield of the indium- and tin-containing sample substrate, so areas with less transmission and thus more electron-scattering biological material appear as darker areas, as first noted by Pluk et al^16^. Moreover, these overlays show that these high contrast areas are not part of the actin fibre but of the FA as identified on the basis of the paxillin fluorescent signal. Such high contrast areas were observed in 98% (95 FAs) of FAs (Table 1). Further examination showed that in almost three quarters of FAs, the high contrasting areas and the associated F-actin fibre are fork-shaped, with the actin as the stem and the high contrasting FA areas as the fork (Fig. 2 and 3). High contrasting areas within the focal adhesion (identified by the presence of paxillin fluorescence) were defined as forked when they were partially intersected by an area of lower contrast splitting the high contrast area into two sides (fork). The angle between the two sides of the fork varied between 7 and 48 degrees, with an average angle of ~20 ± 2.0 degrees (± twice SEM). The average length of the fork was roughly two micrometers which corresponded to ~60 ± 4.5% of the long axis of the FA (Fig. 4 and Table 2). Looking at the two sides of the fork individually, the shortest side was on average ~51 ± 4.8% of the longitudinal FA axis (1.7 ± 0.03 μm) and the longest side ~68 ± 4.8% (2.3 ± 0.03 μm). With regard to symmetry, the average differences between the two sides of the fork was 17.0 ± 3.5% of the FA axis, with a minimum difference of only 0.1% and a maximum difference of 90.3%. Lastly, there was as a trend for longer FAs to have a smaller angle, suggesting the angle of the forks might decrease as FAs mature and grow (Fig. 4b).

**Figure 1.**
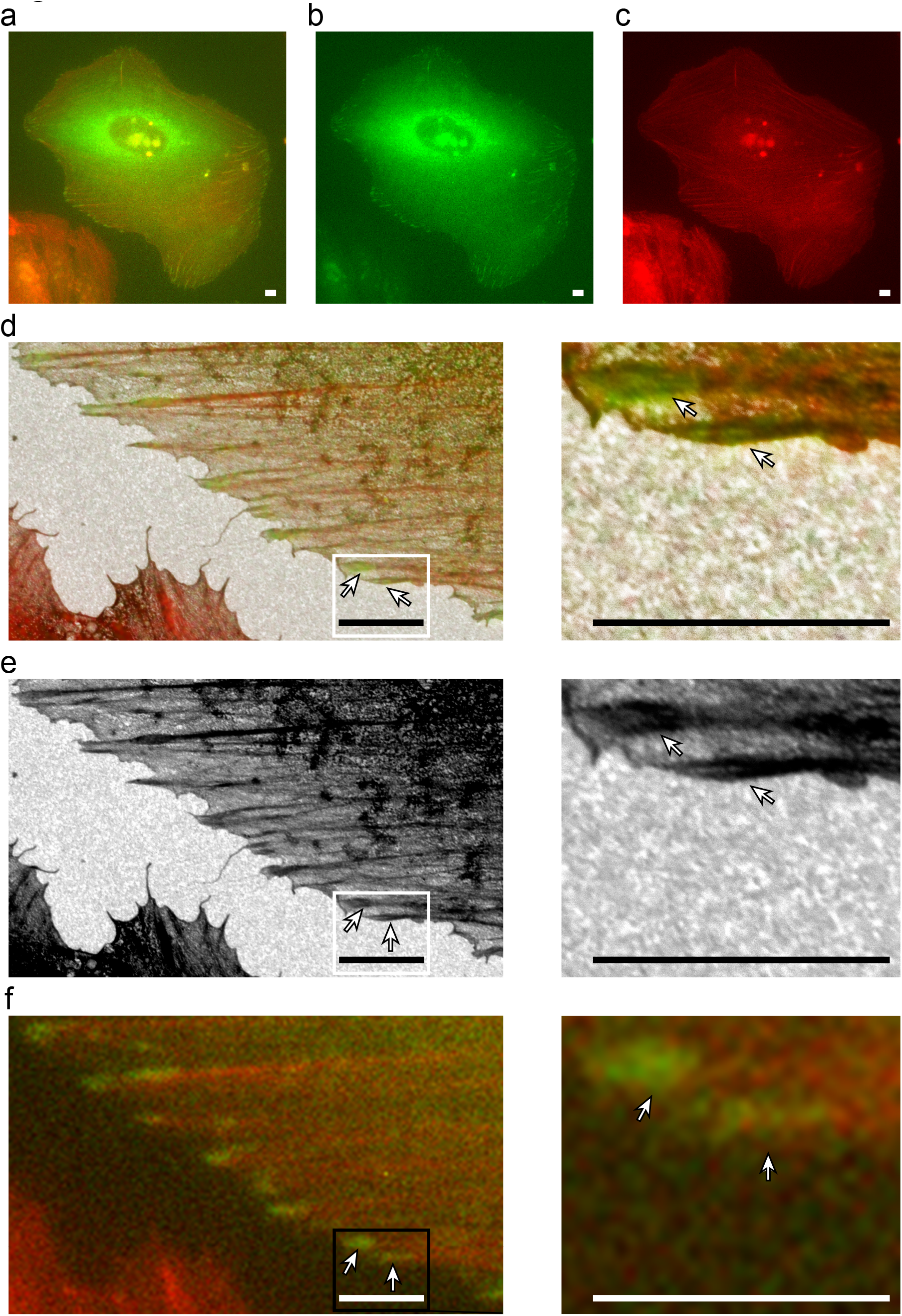
Overlay of fluorescent and SEM images reveals high contrast FA areas around the F-actin entry site. Images of a U2OS cell stably expressing GFP-tagged paxillin and stained with phalloidin cultured on a collagen-coated ITO coverslip. **(a)** Merged image, **(b)** green channel showing FAs where paxillin is localised, **(c)** red channel showing phalloidin stained F-actin. Scalebar: 5 μm. **(d)** Overlay image of a section of the fluorescent image in a with the corresponding SEM image (left), and zoom in of boxed area (right). Arrows indicate FAs with characteristic fork-shapes, where the fork is formed by higher contrast areas of the FA and the stem is formed by the F-actin fibre entering the FA. Scalebar: 5 μm. **(e)** SEM channel of the image in d. **(f)** Fluorescent channel of the image in d.

**Figure 2.**
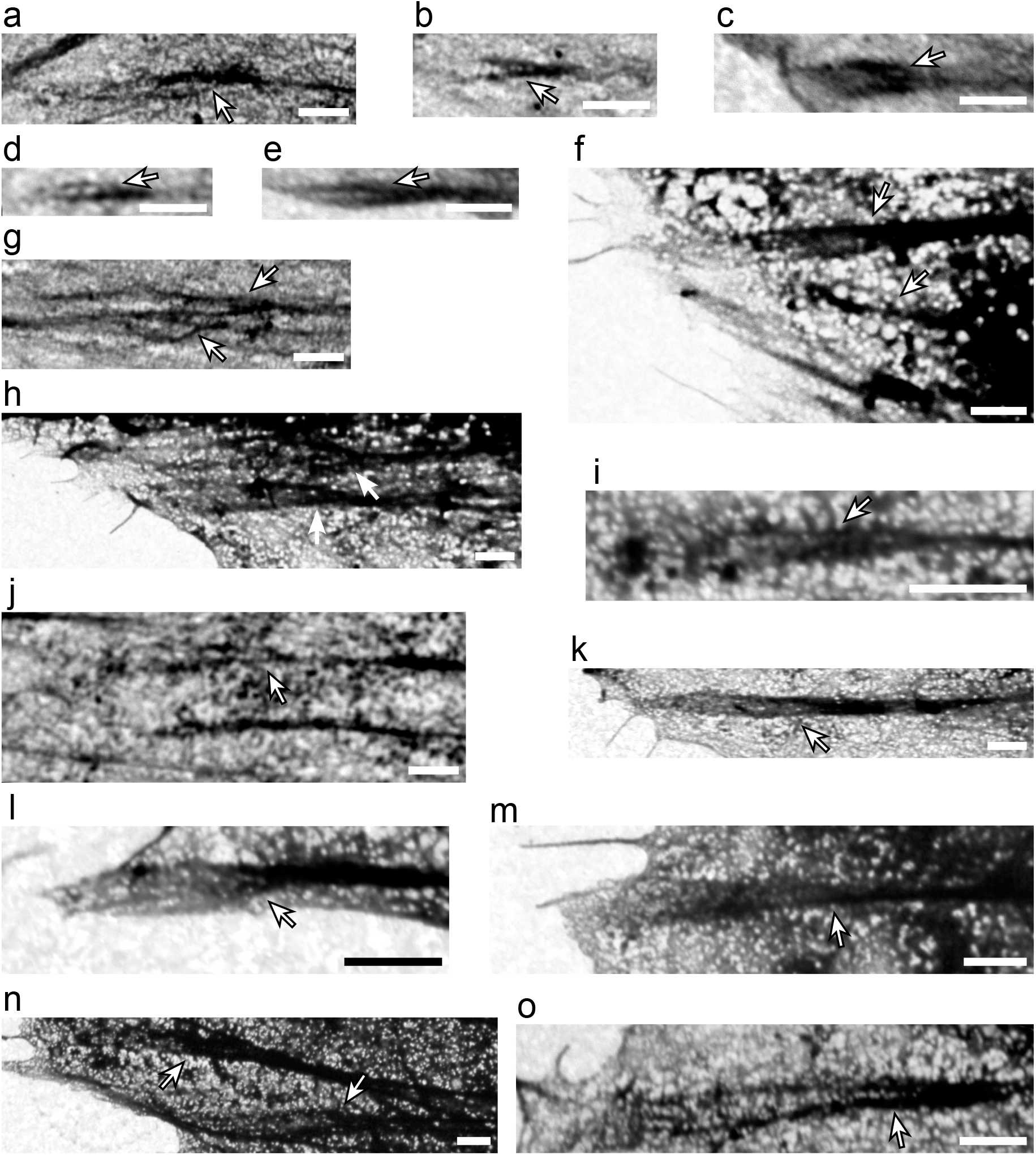
Representative examples of fork-shaped high contrast areas around the F-actin entry side. (**a-o**) High-magnification SEM images of FAs in which the high contrast areas around the F-actin side forms the characteristic fork-like shape (arrows). Scalebar: 1 μm.

**Figure 3.**
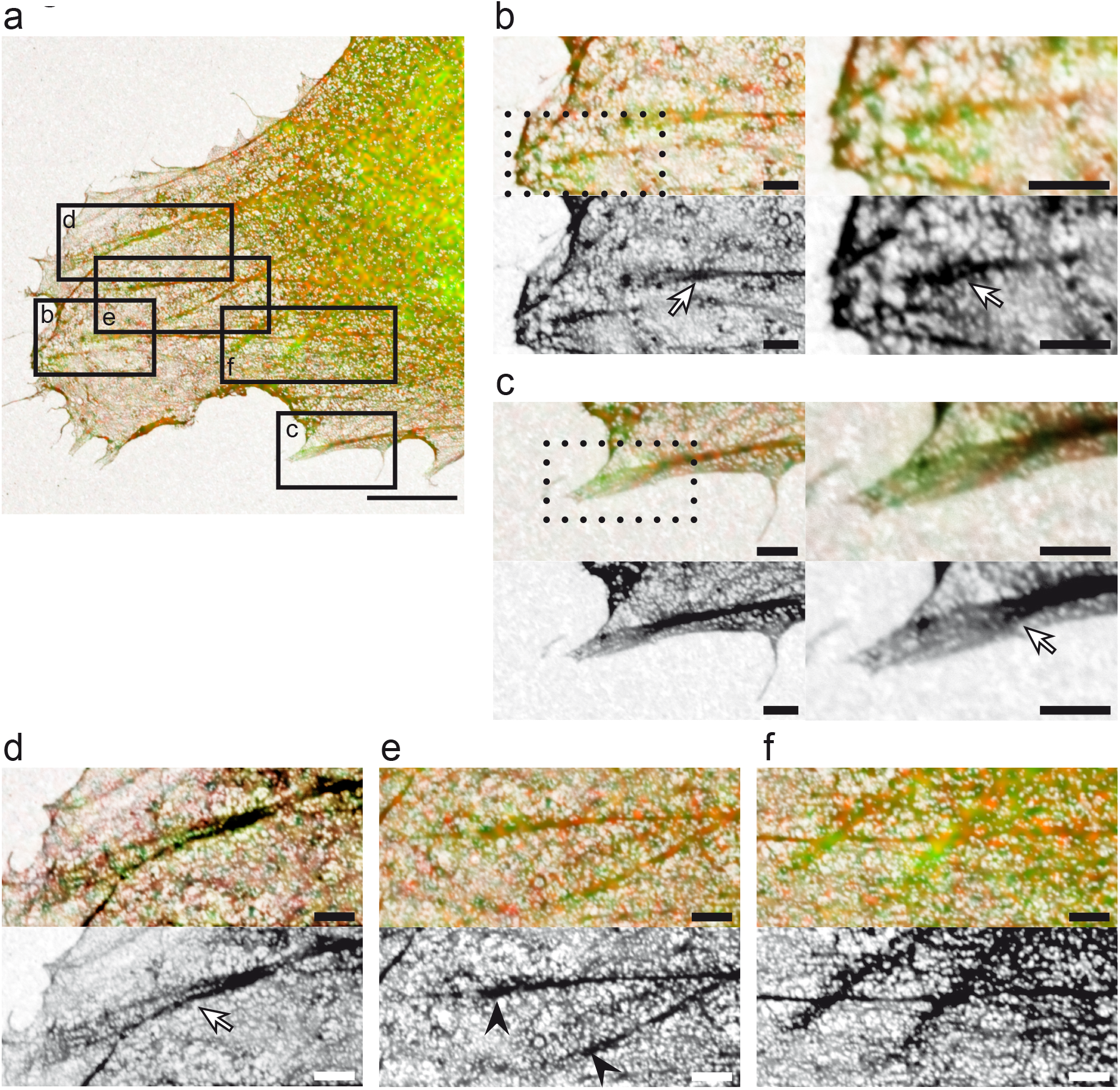
High contrast FA areas around F-actin entry site are a common feature. (**a**) Overlay of SEM image and fluorescent image, with paxillin-stained FAs in green and phalloidin-stained actin in red. Boxes b-f indicate magnifications shown in panels b-f. Scalebar: 5 μm. (**b-e**) Magnifications of corresponding boxed areas in a. Top: overlay image; bottom SEM image. Scalebar: 1 μm. Arrows indicate FAs with characteristic fork-shaped high contrast areas around the F-actin entry site, arrow heads (in e) indicate FAs with high contrast areas not forming a fork-shape. For b and c right images are magnifications of the boxed areas in the left images. (**f**) Magnification of boxed area f in a. Top: overlay image; bottom: SEM image. For the shown FAs the shape of the high contrast areas could not be determined, probably due to the increased thickness of the cell at this more inward location. Scalebar: 1 μm.

**Figure 4.**
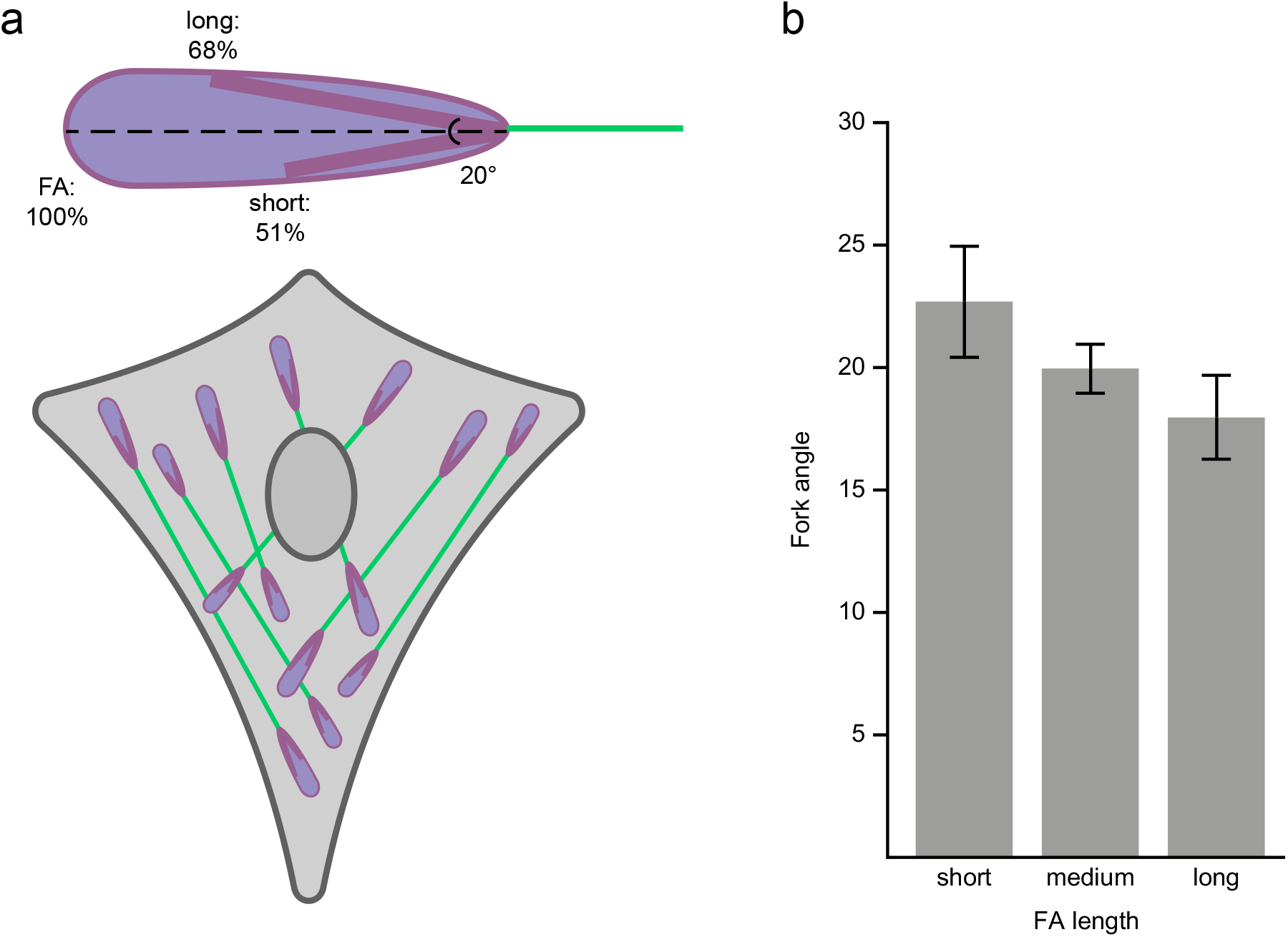
Quantitative measurements of the fork-shape. (**a**) Schematic representation of the average fork-shape observed within FAs based on a quantitative analysis of all observed fork-shapes (see material and methods) (**b**) There is a trend for the angle between the two sides of the fork to be decreased as the length of the FA increases. For this analysis the FAs were split into 3 groups, short: the 24 shortest FAs (ranging from 1.4-2.5 μm), medium: 24 FAs of medium length (ranging from 2.5-3.7 μm) and long: the 25 longest FAs (ranging from 3.7-7.4 μm). Error bars represent SEM.

**Table 1.** The majority of FAs have Fork shaped high contrast areas around the FA entry site. Results of the categorisation of the 97 imaged FAs as independently categorised by two observers. The results for each observer, the percentage of FAs placed in the same category by both observers (% of overlap between observers) and the average results are given.

|  | **FAs with higher contrast at**<br>**actin entry site** | **FAs with fork**<br>**shape** |
| --- | --- | --- |
| **Observer 1** | 97.9% (95 FAs) | 73.2% (71 FAs) |
| **Observer 2** | 97.9% (95 FAs) | 74.2% (72 FAs) |
| **% of overlap**<br>**between observers** | 100% | 94.8% |
| **Average** | 97.9% | 73.7% |

**Table 2.** Quantitative analysis of the fork-shapes. Results of a quantitative analysis of the observed fork shaped areas (n=73). For each FA the longest and shortest side of the fork were measured and their lengths were calculated relative to the longest axis of the FA. In addition the angle between the two sides was measured. For fork symmetry the average difference in length between the long and the short side of the fork relative to the longest axis of the FA was calculated

|  | FA length<br>(long axis;<br>μm) | Fork angle<br>(°) | Fork length<br>(% of FA axis) | Fork symmetry<br>(% of FA axis) | Short side fork (% of FA axis) | Long side fork (% of FA axis) | Fork length (average; μm) | Fork length (short side μm) | Fork length (long side) |
| --- | --- | --- | --- | --- | --- | --- | --- | --- | --- |
| **Average** | 3.30 | 20.2 | 59.7 | 17.0 | 51.1 | 68.2 | 1.98 | 1.69 | 2.26 |
| **Min** | 1.39 | 7.4 | 16.1 | 0.1 | 14.9 | 16.3 | 0.47 | 0.35 | 0.50 |
| **Max** | 7.42 | 47.6 | 90.3 | 65.4 | 88.8 | 99.2 | 5.76 | 5.76 | 5.77 |
| **SD** | 1.33 | 8.7 | 19. | 15.1 | 20.6 | 20.5 | 1.06 | 1.03 | 1.17 |
| SEM | 0.02 | 1.0 | 2.2 | 1.8 | 2.4 | 2.4 | 0.01 | 0.01 | 0.02 |

To our knowledge this is the first report on forked structures with high contrast seen in FAs, possibly because by using correlative fluorescent images, the FA boundaries are clearly defined, whereas based on the SEM images alone the high contrast areas might easily be mistaken for parts of the actin fibre. However, with this new knowledge of exactly what to look for, we were now able to identify (hints of) the fork-shape in previously published EM images of FAs, that were apparently not recognized at that time^10,23,24^. We also note that, while we followed a sample preparation procedure aimed specifically at preservation of fluorescence, these other works used EM-specific sample preparation protocols, indicating the fork-shapes observed here are not an artefact of our fixation procedure.

The nanoarchitecture along the z-axis of FAs has been well described previously^25^. Three different layers have been recognised: (1) the so-called integrin signalling layer (ISL) closest to the adherent membrane (within ~10-20 nm) which includes the cytoplasmic tails of the transmembrane integrin receptors, focal adhesion kinase and paxillin, (2) the actin-regulatory layer (ARL) at the top, where mainly directly actin-binding proteins such as zyxin, vasodilator-stimulated phosphoprotein (VASP) and α-actinin are found, and (3) the force transduction layer (FTL) in between (from ~10-20 to ~50-60 nm from the adherent membrane) of which talin is the most well-known.

The nanoscale structure along the lateral and longitudinal axis of FAs has been investigated to a lesser extent. Along the lateral axis FAs were shown to consist of repeating linear subunits about 300 nm in width^26,27^. Along the longitudinal axis of FAs, different proteins (vinculin, zyxin, paxillin and integrins) were shown to form nanoclusters, suggesting the FA complex as a whole might also be composed of discrete nanoclusters^28–32^. Our data confirms heterogeneity along the longitudinal axis is a feature of the FA complex as a whole, since we are observing the FA complex in an unstained state instead of labelling a specific protein type. More importantly, we provide a location within the FA complex and a specific shape to this heterogeneity, namely a fork at the tip where the actin fibre enters the FA.

On a non-structural level, a few specific parameters have been shown to vary along the longitudinal axis of FAs, for example the binding dynamics of the hallmark FA protein paxillin as was shown in different studies using different approaches^33–36^. Traction forces and molecular tension of FA proteins are also non-uniformly distributed along the longitudinal axis^37,38^. Surprisingly, maximum forces are not found around entry sites of actin fibres, but instead at the opposite tips of the FA. It has been shown that the level of paxillin phosphorylation decreases when force is increased and indeed the level of paxillin phosphorylation has also been demonstrated to vary along FAs^39–41^. This could also be contributing to the formation of the observed high contrast areas, since here the force experienced by the proteins is the lowest, increasing the phosphorylation level of paxillin and perhaps other proteins as well. Strong local increases in the presence of slightly heavier elements than carbon, like phosphor, could further enhance the contrast in SEM images, in addition to a higher density of biological material.

A final example of heterogeneity along the longitudinal axis of FAs involves the activity of vinculin. Vinculin is a large adaptor FA protein like paxillin, but it has a head and a tail domain connected by a flexible linker, allowing vinculin to adopt open (active) and closed (inactive) conformations^42^. This allows assessment of vinculins activity levels through the use of a Foster resonance energy transfer (FRET) biosensor probe, which revealed that FAs at the retracting edge exhibit a gradient of vinculin activity along their lateral axis, with activity increasing towards the actin entry site^43^. A later study was able to generalise these results to FAs beyond the retracting edge^44^. It showed that inactive vinculin associates with the FA layer closest to the adherent membrane (ISL) by binding to phosphorylated paxillin, while talin causes vinculin activation and a shift to the higher FA layers where it binds actin. It was found that inactive vinculin in the ISL is significantly enriched at the FA tip where the actin fibre enters. As inactive vinculin binds to phosphorylated paxillin, this ties in well with studies showing that the forces at actin fibre entry sites are the lowest, leading to higher paxillin phosphorylation levels. The opposite FA tip is significantly enriched in activated vinculin located in the higher FTL and ARL layers, again demonstrating a vinculin activation gradient along FAs. However, our data shows that heterogeneity along the x-axis is a general feature of FAs and extends beyond the activity levels of a single protein like vinculin. Differing levels in one protein alone would not explain the increased contrast in our SEM images around the actin fibre entry site.

Based on the literature, next to paxillin and vinculin, α-actinin is another protein likely to be involved in the formation of the observed forked shape. In a publication using a genetically coded tagged version of α-actinin specifically designed for use with EM, a forked shape analogous to ours can be clearly seen but was not specifically mentioned^24^.

Summarizing, we show that the FA complex is altered around the actin fibre entry site leading to enhanced SEM contrast compared to the rest of the FA. In correlative fluorescence SEM images almost all FAs showed differential levels of contrast and in nearly three quarters of FAs this took the form of fork-shaped structures flanking the actin fibre entry point. Contrast is increased in SEM either when a structure is more dense or when more heavy elements are present, or both. Based on previous literature, it could be hypothesised that proteins likely to be involved in the formation of the fork-shaped structure are paxillin, which is more heavily phosphorylated around the actin entry site, vinculin, the inactive form of which binds to phosphorylated paxillin and is enriched around the entry site, and α-actinin which increases density around the entry site by accumulating here into a fork-shaped structure.

## Materials and Methods

### Cell culture

U2OS cells stably expressing GFP-tagged paxillin were cultured in phenol-red free DMEM (Lonza) at 37°C and 5% CO2. Culture media were supplemented with 10 % FCS (Gibco), 2 mM L-glutamine, 100 U/ml Penicillin, 100 μg/ml Streptomycine and 100 μg/ml G418.

### Sample preparation

For experiments Indium-Tin-Oxide (ITO) coated glass coverslips (22 x 22 x 0.17 mm, Optics Balzers) were coated overnight at 4°C with PureCol bovine collagen type I (Advanced Biomatrix) at a final concentration of 10 μg/ml. Cells were seeded onto coated coverslips and maintained for another 36 h at 37°C and 5% CO2. Cells are chemically fixed with 3.75% paraformaldehyde for 12 minutes and permeabilised with 0.2% Triton-X100 for 10 minutes, before and after each step the cells are washed thrice with PBS. We note that the formaldehyde fixatives penetrate the cells rapidly but may be extracted by repeated washing. We used this procedure in order to preserve the fluorescence in the cells. The more commonly used osmium tetroxide fixate quenches the fluorescence, while glutaraldehyde, which would cross-link more permanently than paraformaldehyde, is auto-fluorescent. As documented in the manuscript, we found evidence for the appearance of the fork-like FA structure in EM images in literature under conditions where stronger fixation was used. Cells were stained with CF405M Phalloidin (Biotium) following the manufacturer’s protocol. Just prior to imaging the samples were dehydrated using an ethanol sequence (2 minutes in 70%, 90%, 95%, and 100% ethanol successively).

### Imaging

Integrated fluorescence microscopy and SEM was carried out using a SECOM integrated microscope (Delmic) retrofitted to a SEM (FEI Verios). The SECOM was equipped with an LED light source (Lumencor Spectra), which was used at 405 nm and 475 nm for Phalloidin and GFP respectively with 140 mW total excitation power. Emission was detected using a multi-band filter (Semrock LED-DA/FI/TR/Cy5-4X-A-000) and a CCD camera (Andor Zyla). Exposure time was typically set at 1 second. The ITO cover slide was mounted to the bottom of a SECOM sample holder ring using carbon tape. An extensive overview of imaging procedure in the integrated microscope was published previously^18^. Fluorescence images were recorded after closing the SEM chamber but before vacuum pump down. After recording the fluorescence images, the SEM was pumped to high vacuum mode and SEM images of selected regions of interest based on fluorescence expression were acquired. The sample was positioned at a working distance of 4.3 mm and back-scattered electrons were detected using a concentric backscatter detector (CBS). Images (4096 x 3775 pixels) were recorded at 2 keV electron energy, a current of 1.6 nA, and 10 μs pixel dwell time. Images in Figures 1, 2a, 2l and 2o were recorded at 3 keV energy and 0.4 nA.

### Analysis

To make overlay images the EM image is opened in black and white as the background layer in Photoshop. The fluorescence image is also opened in Photoshop, scaled to the same size as the SEM image (based on the known pixel dimensions) and placed as a separate layer in colour on top of the SEM background layer. The precise overlay is done by hand, mainly based on the F-actin network which due to phalloidin staining is clearly visible in both imaging dualities. To determine the prevalence of fork shaped high contrast areas (defined as any high contrast area that is partially intersected by an area of lower contrast, splitting the high contrast area into two sides) in the FAs of our data set all FAs were assigned a number. In total 122 FAs were counted based on the fluorescent images and of these 97 were also clear enough in the SEM image to be able to determine the shape of the high contrast areas. The reasons to classify an FA as not clear enough in the SEM image to be able to determine the shape of the high contrast areas were in order of occurrence 1) the FA lies too far away from the edge of the cell where the cell is thin enough to image through with the SEM so the FA is lost in the signal coming from the cytoplasm/cytoskeleton of the cell 2) the FA is too small in the SEM image so while it is possible to make out the FA in the SEM image the resolution of the SEM image is not sufficient to allow the shape of the high contrast area to be determined or 3) the FA itself or the connecting F-actin fibre is not visible in the SEM image which doesn’t allow anything to be said about the shape of a putative darker area at the place where the actin connects to the FA as this connection point is not visible. Of these 97 FAs two people independently counted the number of high contrast areas and the number of high contrast areas with a Y-shape, with very comparable results: 95 FAs have a high contrast area and for 71/72 FAs this area is fork shaped, we reported the average. To allow a more quantitative analysis of these fork shapes, all fork shapes and the longest axis of the FAs were also traced by hand using the Fiji inbuilt angle and line-drawing and measuring tools. Care was taken that for the angle measurements using the angle measurement tool only the beginning of the fork shape was used.

